# An ancestral SARS-CoV-2 vaccine induces anti-Omicron variants antibodies by hypermutation

**DOI:** 10.1101/2023.03.15.532728

**Authors:** Seoryeong Park, Jaewon Choi, Yonghee Lee, Jinsung Noh, Namphil Kim, JinAh Lee, Geummi Cho, Sujeong Kim, Duck Kyun Yoo, Chang Kyung Kang, Pyoeng Gyun Choe, Nam Joong Kim, Wan Beom Park, Seungtaek Kim, Myoung-don Oh, Sunghoon Kwon, Junho Chung

**Author notes:** These authors contributed equally to this work (Seoryeong Park, Jaewon Choi). **Corresponding author** S. Kim, M. Oh, S. Kwon or J. Chung.

## Abstract

The immune escape of Omicron variants significantly subsides by the third dose of an mRNA vaccine. However, it is unclear how Omicron variant-neutralizing antibodies develop under repeated vaccination. We analyzed blood samples from 41 BNT162b2 vaccinees following the course of three injections and analyzed their B-cell receptor (BCR) repertoires at six time points in total. The concomitant reactivity to both ancestral and Omicron receptor-binding domain (RBD) was achieved by a limited number of BCR clonotypes depending on the accumulation of somatic hypermutation (SHM) after the third dose. Our findings suggest that SHM accumulation in the BCR space to broaden its specificity for unseen antigens is a counter protective mechanism against virus variant immune escape.

## Main

Since the emergence of severe acute respiratory syndrome coronavirus 2 (SARS-CoV-2), over 14 million sequences of variants have been collected and shared via the Global Initiative on Sharing All Influenza Data (GISAID) ^1^. While most mutations have little effect or are detrimental to the virus, a small subset of mutations may provide a selective advantage leading to a higher reproductive rate^2^. The spike protein, a viral coat protein that mediates viral attachment to host cells and fusion between the virus and the cell membrane, is the primary target of neutralizing antibodies^3^. Serological analysis has shown that the receptor-binding domain (RBD) of the spike protein is the target of 90% of neutralizing activity in the immune sera^4, 5^. In this context, the RBD has become the essential component of most mRNA-, adenovirus-, and recombinant protein-based vaccines^6^. However, Omicron variant BA.1 has accumulated 15 mutations in RBD^7^, resulting in a 22-fold reduction in neutralization by plasma from vaccinees receiving two doses of the BNT162b2 vaccine^8^. This has led to reduced efficacy of most commercially available monoclonal antibodies and antibodies under development against BA.1^7, 9^. Although bivalent vaccines have been developed to overcome the immune evasion of Omicron variant^10–13^, the majority of the population has received only monovalent vaccines to date. Fortunately, it has been proven that a third dose of the BNT162b2 monovalent vaccine can neutralize BA.1^14–18^. However, the mechanism by which Omicron variant-neutralizing antibodies are generated through repeated exposure to the ancestral spike protein remains unclear. In this study, we analyzed the chronological B-cell receptor (BCR) repertoires of BNT162b2 vaccinees and traced the development of Omicron variant-neutralizing antibodies.

### Vaccination and plasma antibody levels

We followed 41 healthcare workers from Seoul National University Hospital who received two doses of the BNT162b2 mRNA vaccine with a three-week interval and a third dose at approximately nine months after the first dose (Fig. 1a). Peripheral blood samples were collected six times: once before the first dose, three times after each dose, and two times between the second and third doses. The vaccinees did not report a history of COVID-19 infection, and the absence of plasma IgG against the nucleocapsid (N) protein of SARS-CoV-2 was confirmed in all vaccinees (Extended Data Fig. 1). In an enzyme-linked immunosorbent assay (ELISA), plasma levels of IgA and IgG against the ancestral RBD were significantly increased after the second dose (Fig. 1b and Extended Data Fig. 2). However, when vaccination ELISA was performed against BA.1 RBD, a third dose was essential to achieve elevated levels of IgA and IgG, which also reacted to the RBD of Omicron subvariant BQ.1.1 (Fig. 1b, Extended Data Figs. 2 and 3). This observation was in accord with prior reports that neutralizing activity of plasma against the Omicron variant was detectable only after the third dose^19, 20^, while the second dose could induce Omicron variant-neutralizing memory B cells^21^.

**Fig. 1.**
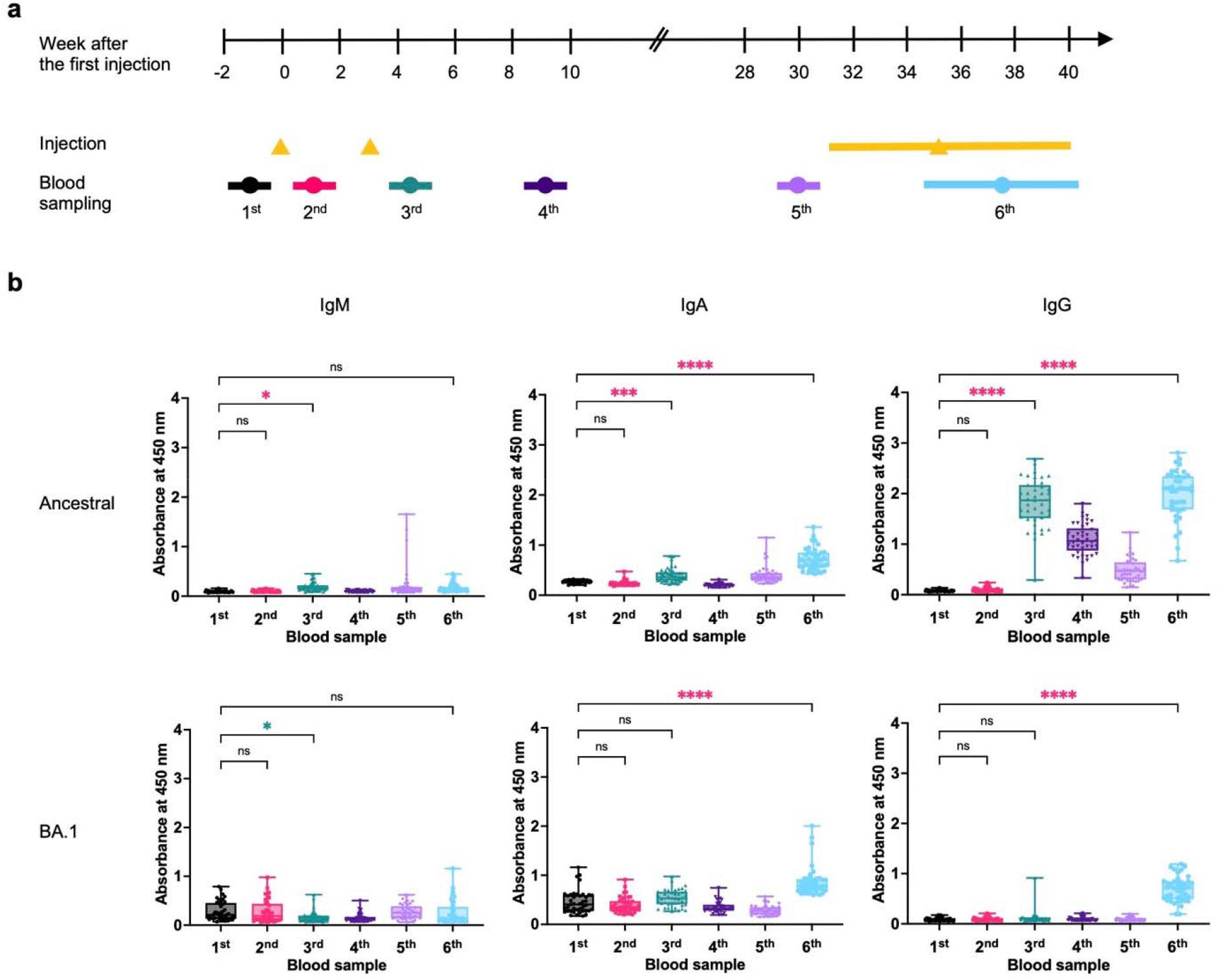
Vaccination, blood sampling and plasma antibody levels. **a,** Vaccinees received the first two doses at a three-week interval and the third dose between the 31st and 39th weeks after the first dose. Blood was collected six times: pre-vaccination (1^st^); 1 week after the first dose (2^nd^); 1, 6, and 30 weeks after the second dose (3^rd^, 4^th^, and 5^th^); and one to four weeks after the third dose (6^th^). **b,** Plasma levels of antibodies against SARS-CoV-2 ancestral and BA.1 RBDs were measured after 2,500-fold dilution. All experiments were performed in duplicate, and the average value for each vaccinee was plotted. The P value was calculated using one-way ANOVA. ns, not significant, p > 0.05; *, p < 0.05; ***, p < 0.001; ****, p < 0.0001. The increases and decreases in antibody levels are marked with red and green asterisks, respectively.

### Selection and characterization of BA.1 RBD-binding antibody clones

We selected six vaccinees and generated a single-chain variable fragment (scFv) phage display library using cDNA prepared from the sixth blood sample. For the selection of these vaccinees, we reconstituted the *in silico* chronological BCR repertoires of vaccinees by next-generation sequencing (Supplementary Table 1) and compared the frequency of BCR heavy chain (HC) clonotypes listed in the CoV-Ab Dab^22^ database binding the SARS-CoV-2 spike protein. Vaccinee 32, 35, 39 and 43 and vaccine 22 and 27 showed a higher frequency of clonotypes binding to the BA.1 and ancestral spike proteins, respectively, and were selected. In this study, the BCR HC clonotype was defined as a set of immunoglobulin heavy chain sequences that exhibit the same immunoglobulin heavy variable (IGHV) and immunoglobulin heavy joining (IGHJ) genes and homologous heavy chain complementarity-determining region 3 (HCDR3) amino acid sequences with a minimum sequence identity of 80% to a reference sequence. From six libraries, nine BA.1 RBD-binding scFv clones were selected based on biopanning and phage ELISA. A cluster of scFv clones composed of 27-60 clones and 51 other clones sharing IGHV3-53/3-66, IGHJ6, and the same HCDR sequences at the amino acid level was selected from five libraries (Table 1 and Supplementary Table 2). We have previously reported that an IGHV3-53/3-66 and IGHJ6 BCR HC clonotype is present not only in the majority of convalescent patients from ancestral strains but also in normal human populations not exposed to SARS-CoV-2 and can neutralize the virus^23^. The frequent use of the IGHV3-53/3-66 and IGHJ6 gene pair in SARS-CoV-2 neutralizing antibodies was further confirmed by a subsequent report^24–26^. Six scFv clones carrying the IGHV1-69 gene were selected from two libraries, which frequently encoded RBD-reactive antibodies found in convalescent patients from both the ancestral strain and BA.1 variant^27^. Two additional scFv clones were selected from the two libraries.

**Table 1.**
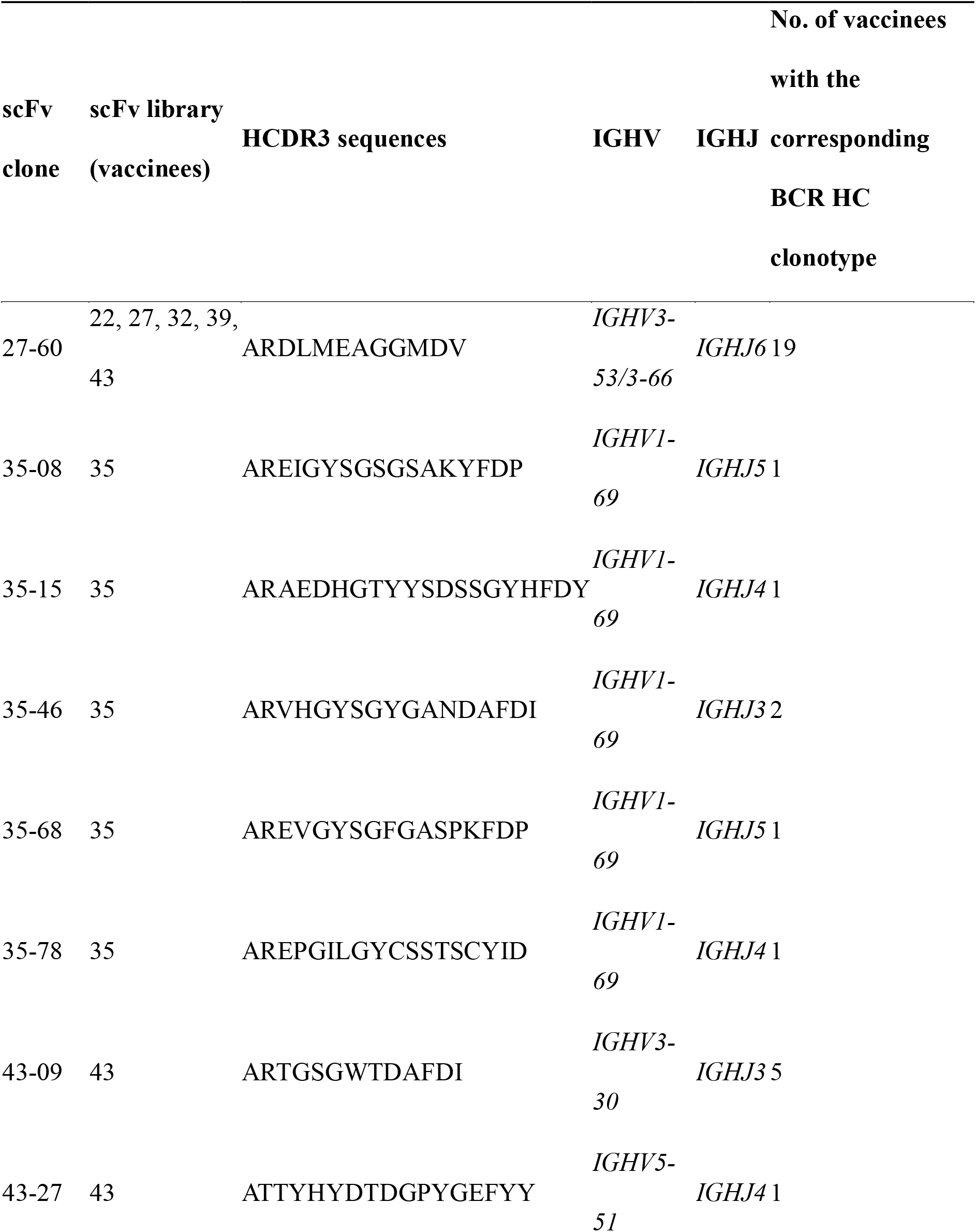
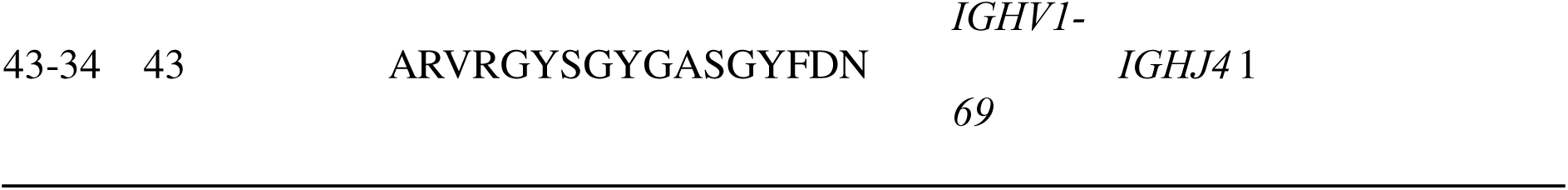
Omicron RBD-reactive scFv clones and their BCR-HC clonotypes

In the recombinant scFv-human Fc-hemagglutinin (HA) fusion protein format, seven scFv clones showed reactivity against the RBD of the ancestral virus, Alpha, Beta, Gamma, and Delta variants and Omicron-sublineage variants (BA.1, BA.2, BA.4/5 and BQ.1.1) (Extended Data Fig. 4a), with half-maximal effective concentrations (EC50) below 350 pM. The 27-60 clone and 35-15 clone showed dramatically reduced reactivity to BA.4/5 and BQ.1.1 RBDs and BQ.1.1 RBD, respectively. In a microneutralization assay, all clones neutralized the ancestral SARS-CoV-2 strain with a half-maximal inhibition concentration (IC50) range of 0.44 to 10.03 μg/ml (Extended Data Fig. 4b). Four clones (35-8, 35-46, 35-68, and 35-78) showed slightly decreased neutralizing activity against the BA.1 variant with an IC50 below 15 μg/ml, while the activity of others was increased to 25.79 to 74.12 μg/ml. Three clones (35-8, 35-68, and 35-78) could also neutralize Alpha, Beta, Gamma and Delta variants with less IC50 values under 12.12 ug/ml.

### Omicron RBD-reactive clonotypes in BCR HC repertoires

BCR HC clonotypes of Omicron RBD-binding clones (Omicron RBD-reactive BCR HC clonotype) were mapped to 293 sequences in the repertoire of 23 vaccinees (Table 1 and Supplementary Table 3). The most frequently identified BCR HC clonotype was 27-60, found in 19 vaccinees (46%), followed clonotypes by 43-09 and 35-46, found in five and two vaccinees (12% and 5%), respectively (Supplementary Table 4). Six BCR HC clonotypes were found only in one vaccinee. Five BCR HC clonotypes, including 27-60, were not found in vaccinees in whom the antibody clones were found. This type of discrepancy is expected to inevitably originate from the incomplete *in silico* reconstitution of the BCR HC repertoire due to the limitation of the throughput of next-generation sequencing^28^. In this regard, the BCR HC clonotypes are expected to be present in a larger population than we observed herein.

Among the 293 BCR HC sequences, 57 and 216 BCR HC sequences were present in the third and sixth repertoires, respectively. Only 20 BCR HC sequences were found in other chronological BCR repertoires. This skewed distribution originated from the expansion and proliferation of B cells encoding the Omicron RBD-reactive BCR HC clonotype immediately after the second and third doses of the vaccine. Thus, we limited our statistical analysis to BCR HC sequences present in the third and sixth repertoires. Rapid class switch recombination (CSR) following vaccination was evident, as the main immunoglobulin subtype of Omicron RBD-reactive BCR HC sequences was IgG_1_ in the third repertoire, followed by IgG_2_, which was maintained in the sixth repertoire (Supplementary Table 4 and Extended Data Fig. 5a). This rapid CSR resulting in the dominance of the IgG_1_ subtype among RBD-reactive BCR HC sequences was also observed in convalescent patients infected with the ancestral virus^23^.

Additionally, somatic hypermutation (SHM) of Omicron RBD-reactive BCR HC clonotypes occurred mainly after the third dose. The average number of SHMs in the IGHV gene of Omicron RBD-reactive BCR HC clonotypes in the sixth repertoire was dramatically increased compared to that in the third repertoire with statistical significance (Extended Data Fig. 5b). The diversity of the HCDR3 sequence also increased from the third repertoire to the sixth repertoire (Extended Data Fig. 5c). Five Omicron RBD-reactive BCR HC clonotypes were mapped to eight vaccinees in both the third and sixth repertoires (Extended Data Fig. 5d). The increase in the average number of SHMs was evident, and statistical significance was confirmed in three vaccinees with the 27-60 clonotype and one vaccinee with the 43-34 clonotype.

### Development of concomitant reactivity to ancestral and BA.1 RBDs via SHM

Clone 27-60 was encoded by IGHV3-53/IGHV3-66 (Fig. 2a), in which CDR1 and CDR2 are known to have two key motifs providing reactivity to ancestral RBD^23, 29^. The BCR HC of clone 27-60 had two SHMs in its IGHV gene sequence, and when we back-mutated the V27 residue in CDR1 or the F58 residue adjacent to CDR2 to the germline sequence, its affinity for the ancestral and BA.1 RBDs was decreased 8.7-fold and greatly diminished, respectively (Fig. 2b). This observation showed that the reactivity of clone 27-60 toward the BA.1 RBD depended on the SHM to a greater degree than its reactivity to the ancestral RBD.

**Fig. 2.**
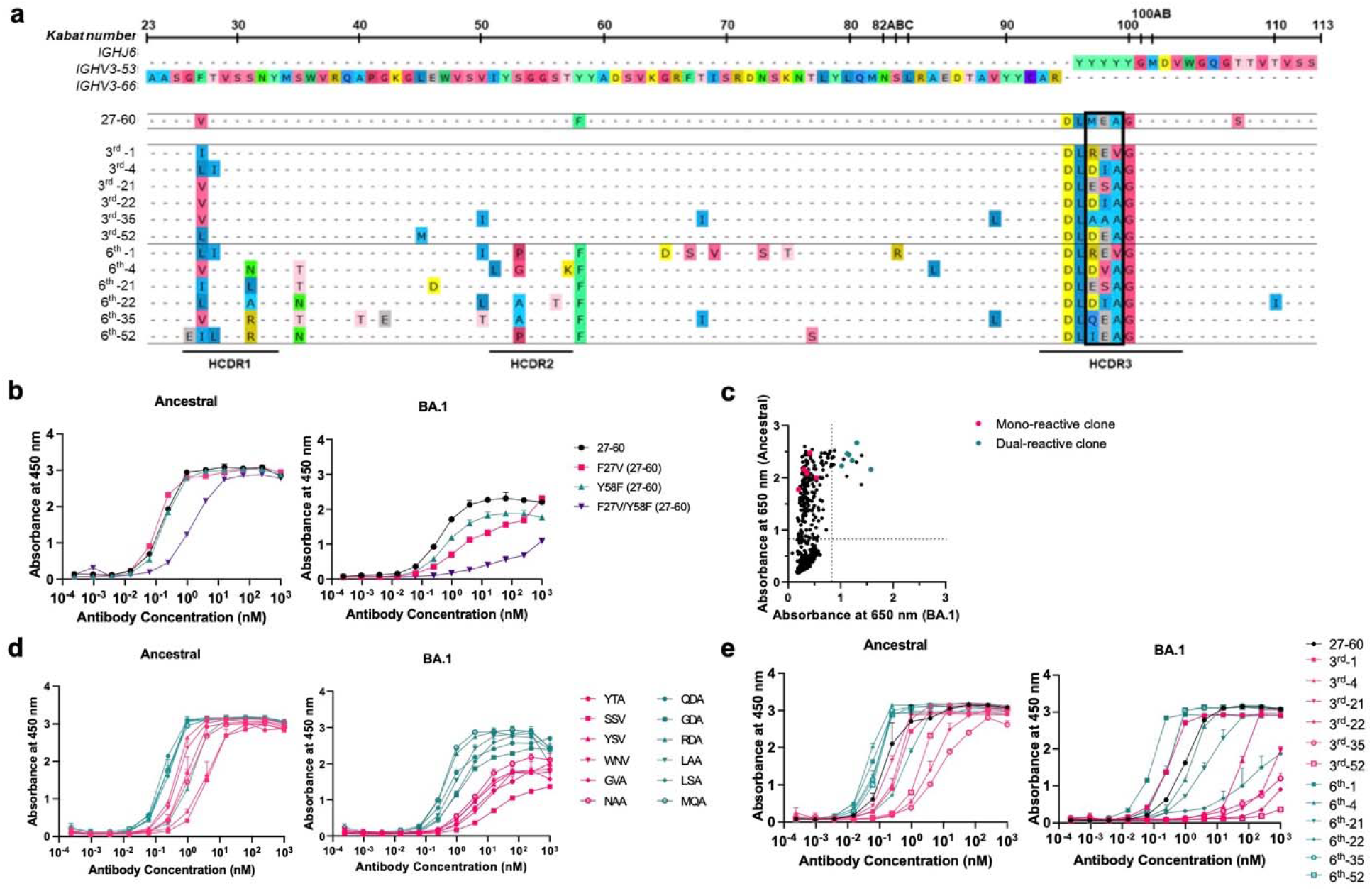
Characterization of the 27-60 BCR HC clonotype. **a,** BCR HC sequences found in six vaccinees in their third and sixth BCR repertoires with the highest frequency. **b,** Reactivity of the recombinant 27-60 scFv-hFc-HA protein and its clones back mutated to the germline sequence. **c,** Reactivity of scFv-displaying phage clones from the HCDR3-randomized library to the ancestral and BA.1 RBDs. The clones labeled in red and green were selected for expression as recombinant scFv-hFc-HA protein. **d,** Reactivity of recombinant scFv fusion proteins of HCDR3-randomized (ARDLXX(A/V)GGMDV) clones to the ancestral and BA.1 RBDs. **e,** Reactivity of recombinant scFv-hFc-HA proteins encoded by individual BCR HC sequences and the light chain gene of clone 27-60 to the ancestral and BA.1 RBDs.

Thereafter, we selected the most frequent BCR HC sequences of the 27-60 clonotype in the third and sixth BCR repertoires of six vaccinees and analyzed their sequences (Fig. 2a). All the BCR HC sequences found in the sixth repertoires had more SHMs than their pairs in the third repertoires. All six BCR HC sequences in the sixth repertoire gained the Y58F mutation, increasing the affinity of IGHV3-53/IGHV3-66 antibodies for the ancestral RBD^30, 31^. There was also diversity in their HCDR3 sequences limited to a motif of three amino acid residues. As the HCDR3 sequence can diversify either at the stage of germline VDJ gene recombination by P- and N-nucleotide addition/deletion or later by SHM, it was not possible to determine the germline HCDR3 sequence of clones 27-60. To analyze the influence of the HCDR3 sequence on reactivity to ancestral and BA.1 RBDs, we generated a scFv phage display with the randomized HCDR3 sequence of ARDLXX(A/V)GGMDV, arbitrarily selected 672 phage clones and checked their reactivity to both RBDs. For the ancestral RBD, 200 phage clones (30%) showed positive reactivity (Fig. 2c, Extended Data Fig. 6 and Supplementary Table 5). However, for the BA.1 RBD, only 18 phage clones (3%) were reactive. All 18 clones were dual-reactive to both RBDs. Six mono-reactive and six dual-reactive phage clones were arbitrarily selected for expression as recombinant scFv-hFc-HA fusion proteins and tested for their affinity for both RBDs. The EC50 values of mono-reactive scFv proteins to the ancestral and BA.1 RBDs were in the ranges of 0.84–3.80 nM and 3.83–17.25 nM, respectively. In addition, dual-reactive scFv proteins showed EC50 values of 0.16–1.29 nM and 0.35–1.26 nM for the ancestral and BA.1 RBDs, respectively (Fig. 2d). These observations proved that it is relatively rare for ancestral RBD-reactive BCR HC clonotypes encoded by the IGHV3-53/3-66 and IGHJ6 genes with the HCDR3 sequence of ARDLXX(A/V)GGMDV (BCR HC XXA/V clonotypes) to develop increased compatible reactivity to the BA.1 RBD. This dual reactivity is likely to be achieved among the large population of antigen-stimulated and proliferating B cells through SHM, rather than in a small number of B cells in the stage of germline VDJ gene rearrangement. Indeed, only one BCR HC XXA/V sequence was found in only one vaccinee in the pre-immune repertoire (Extended Data Fig. 7 and Supplementary Table 6). After the first, second, and third injections, 3, 58, and 138 BCR HC XXA/V sequences were found in two, ten and sixteen vaccinees, respectively. The average frequency of BCR HC XXA/V sequences also increased from 6.06×10^-6^ under pre-immune status to 2.85×10^-5^, 1.91×10^-5^, and 2.41×10^-5^ after the first, second, and third doses, respectively.

To check the effect of SHM accumulation on the affinity for RBDs in 27-60 BCR HC clonotypes, six pairs of BCR HC sequences were coupled with the light chain gene of clone 27-60 (Supplementary Table 7) and expressed as recombinant scFv-hFc-HA fusion proteins. Then, their affinity for the ancestral RBD and BA.1 RBD was determined by ELISA. For the ancestral RBD, all six scFvs of the third repertoire showed notable affinities, with EC50 values less than 7.5 nM. However, only one scFv from vaccinee 1 showed comparable affinity for BA.1 RBD (Fig. 2e and Supplementary Table 8). In the sixth repertoire, four additional scFvs showed affinity for the BA.1 RBD, with EC50 values less than 4.5 nM.

We also analyzed the BCR HC sequences of four additional clonotypes (35-15, 35-46, 43-09 and 43-34) found in both the third and sixth repertoires of identical vaccinees (Extended Data Fig. 8a). All BCR HC sequences in the sixth repertoires showed more SHMs than their paired sequences in the third repertoires. Then, we prepared eight recombinant scFv-hFc-HA fusion proteins in pairs with the light chains of four scFv phage clones. In ELISA, all scFvs from the third repertoire showed a remarkable level of affinity for the ancestral RBD, with EC50 values less than 200 pM, which were well maintained within a 2-fold difference in their paired sequences in the sixth repertoire (Extended Data Fig. 8b). It was exceptional that the BCR HC sequences of the 35-46 clonotype in the third library were encoded by IGHV1-69/IGHJ3 genes without any SHM and showed a significantly high affinity for BA.1 RBD. In the 35-15, 43-09 and 43-34 BCR HC clonotype pairs, the affinity for BA.1 RBD was significantly increased from 0.59, 1.12 and 7.91 nM in the third repertoire to 0.13, 0.09 and 0.14 nM in the sixth repertoire, respectively.

### Expansion of the BCR HC repertoire to RBDs of Omicron subvariants by SHM

We studied how the diversification of the BCR HC repertoire by SHM expands its reactivity to RBDs of Omicron subvariants. We first searched the BCR HC sequences of the 27-60, 35-15, 35-46, 43-09 and 43-34 clonotypes in the third repertoire and found them in nine, one, one, one, and one of the vaccinees, respectively (Supplementary Table 4). Then, we selected the BCR HC sequences with the highest frequency in each vaccinee and used them to construct a new set of BCR HC clonotypes (designated clonotypes-3^rd^) confined to each vaccinee. The greatest numbers of BCR HC sequences in the 27-60, 35-15, 35-46, 43-09 and 43-34 clonotypes-3^rd^ were found in vaccinees 35, 43, 35, 23, and 43, consisting of 86, 8, 24, 2, and 18 sequences, respectively. Based on their HCDR3 sequences, these sequences in the clonotypes-3rd were subdivided into 11, 4, 11, 2, and 9 clusters, respectively. From each cluster, the BCR HC sequences with the highest frequency that were present in the third and sixth BCR HC repertoires were selected, and their genes were synthesized (Extended Data Fig. 9, 10 and Supplementary Table 7, 9). As expected, all the scFv clones in the sixth BCR HC repertoires showed more SHM than those in the third repertoires (Fig. 3), and some of them showed dramatically enhanced binding toward specific types of Omicron variant RBDs. Two scFv clones (35-15/6th-43 and 35-46/6th-35-2) showed potent reactivity confined to either BA.1 or XBB.1.5 RBD with an EC50 below 0.100 nM. The other two scFv clones (35-46/6th-35-1 or 43-09/6th-23) showed the equivalent EC50 values toward RBDs of Omicron subvariants (either BA.1 or BQ.1.1 in addition to XBB.1.5, individually). Another two scFv clones (43-34/6th-43-1 and 43-34/6th-43-3) reacted with the RBDs of all Omicron subvariants tested with same potency. These findings suggest that the accumulation of SHM after the third dose of ancestral SARS-CoV-2 vaccine generated clones with divergent mosaic pattern of specificity toward three major Omicron subvariants. More interestingly, clones reactive to BQ.1.1 and XBB.1.5 achieved their specificity much earlier than their emergence.

**Fig. 3.**
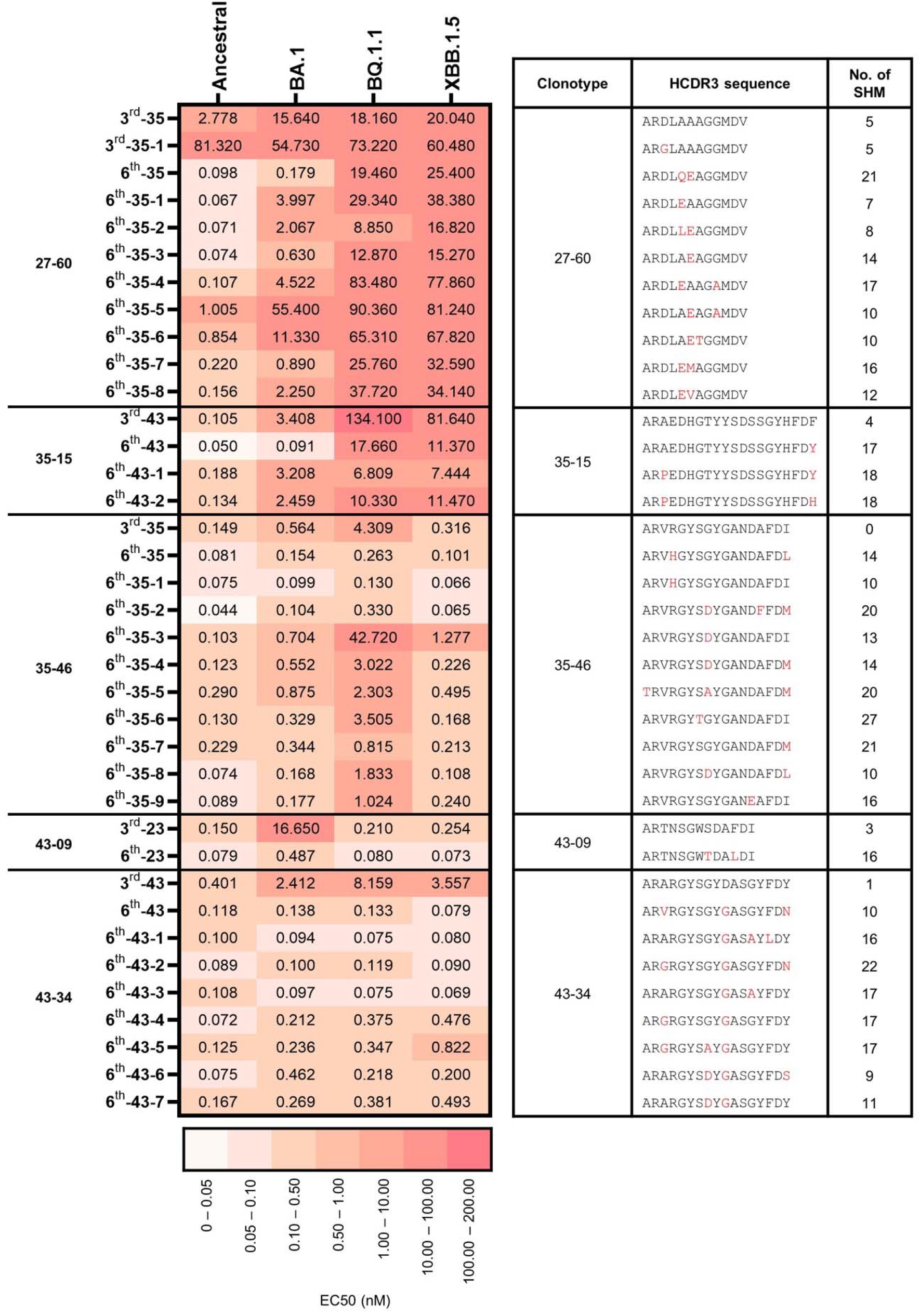
The reactivity of the scFv clones of HCDR3-based clusters in the 27-60, 35-15, 35-46, 43-09, and 43-34 BCR HC clonotypes-3^rd^. The EC50 values to Ancestral, BA.1, BQ.1.1, and XBB.1.5 RBDs of the scFv clones with the highest frequencies of HCDR3-based clusters in each clonotype-3^rd^ are displayed with their HCDR3 sequences and the quantities of SHM.

Our study showed that the third dose of the mRNA vaccine encoding the ancestral viral spike protein induced the accumulation of SHM in ancestral RBD-reactive antibody clones and generated a panel of BCRs with additional divergent reactivity to RBDs of Omicron subvariants, which may contribute to the dramatically increased plasma level of Omicron RBD–reactive antibodies and may have the potential to counteract novel SARS-CoV-2 variants yet to emerge. We believe that this expansion in the specificity of the BCR repertoire to unseen antigens via SHM is a protective immune mechanism that evolved in response to the challenge of viral immune escape.

## Methods

### Study participants

Peripheral blood sampling was approved by the Institutional Ethics Review Board of Seoul National University Hospital (IRB approval number, 2102-032-1193). The vaccinees had a median age of 30 years (range 23-62) and showed a nearly equal distribution of males and females (46% and 54%, respectively). Peripheral blood mononuclear cells (PBMCs) and plasma were separated using Lymphoprep (STEMCELL) Ficoll (Cytiva) following the manufacturer’s instructions. Total RNA was isolated using TRIzol Reagent (Invitrogen) according to the manufacturer’s instructions.

### ELISA

Plasma and phage ELISAs were performed in 96-well microtiter plates (Corning, 3690) coated with 100 ng of recombinant SARS-CoV-2 proteins (Sino Biological, Ancestral RBD, 40592-V08H; Alpha RBD, 40592-V08H82; Beta RBD, 40592-V08H85; Gamma RBD, 40592-V08H86; Delta RBD, 40592-V08H90; Omicron BA.1 RBD, 40592-V08H121; Omicron BA.2 RBD, 40592-V08H123, Omicron BA.4/5 RBD, 40592-V08H130; Omicron BQ.1.1 RBD, 40592-V08H143; Ancestral N, 40588-V08B) in coating buffer (0.1 M sodium bicarbonate, pH 8.6) as described previously ^23^. Briefly, the plates were coated with the antigen by incubation at 4°C overnight and blocked with 3% bovine serum albumin (BSA) in PBS for 1 hour at 37°C. Then, serially diluted serum (100-, 500-, 2,500-fold) or phage supernatant (twofold) in blocking buffer was added to the wells of microtiter plates, followed by incubation for 1 hour at 37°C. Then, the plates were washed three times with 0.05% PBST. Horseradish peroxidase (HRP)-conjugated goat anti-human IgM and IgA (Invitrogen, A18835 and A18781, 1:5,000), rabbit anti-human IgG antibody (Invitrogen, 31423, 1:20,000) and HRP-conjugated anti-M13 antibody (Sino Biological, 11973-MM05T-H, 1:4,000) were used to determine the amount of bound antibody or M13 bacteriophage. A 3,3′,5,5′-tetramethylbenzidine liquid substrate solution (Thermo Fisher Scientific Inc.) was used as an HRP substrate, and the absorbance was measured at 450 or 650 nm using a microplate spectrophotometer (Thermo Fisher Scientific Inc., Multiskan GO) depending on the use of 2 M sulfuric acid as the stop solution. All assays were performed in duplicate.

In ELISA for the recombinant scFv-hFc-HA fusion proteins, the wells of microtiter plates (Corning) were first coated with 100 ng of mouse anti-His antibody (Invitrogen, MA1-21315) and blocked. Then, the recombinant SARS-CoV-2 RBD protein with a polyhistidine tag (100 nM) was added to the wells. After brief washing, the scFv-hFC-HA fusion proteins were serially diluted fourfold from 1 μM to 0.24 pM in blocking buffer and added to wells of a microtiter plate. HRP-conjugated rat anti-HA antibody (Roche, 12013819001, 1:1,000) was used to determine the amount of bound antibody. All assays were performed in duplicate.

### Next-generation sequencing

Genes encoding the variable domain of the heavy chain (V_H_) and part of the first constant domain of the heavy chain (CH1) domain (V_H_-CH1) were amplified using specific primers, as described previously^23^. All primers used are listed in Supplementary Table 10. Briefly, total RNA was used as a template to synthesize complementary DNA (cDNA) using the SuperScript IV First-Strand Synthesis System (Invitrogen) with specific primers (primer No. 1–8) targeting the CH1 domain of each isotype (IgM, IgD, IgG, IgA, and IgE) according to the manufacturer’s protocol. After cDNA synthesis, 1.8 volumes of SPRI beads (Beckman Coulter, AMPure XP) were used to purify cDNA, which was eluted in 35 μl of water. The purified cDNA (15 μl) was subjected to second-strand synthesis in a 25 μl reaction volume using IGHV gene–specific primers (primer No.9–14) and a KAPA Biosystems kit (Roche, KAPA HiFi HotStart). The PCR conditions were as follows: 95°C for 3 min, 98°C for 30 s, 60°C for 45 s, and 72°C for 6 min. After second-strand synthesis, dsDNA was purified using 1 volume of SPRI beads, as described above. V_H_-CH1 genes were amplified using purified dsDNA (15 µl) in a 25 μl reaction volume using primers containing indexing sequences (primers 15 and 16) and a KAPA Biosystems kit. The PCR conditions were as follows: 95°C for 3 min; 25 cycles of 98°C for 30 s, 60°C for 30 s, and 72°C for 1 min; and 72°C for 5 min. PCR products were subjected to electrophoresis on a 1.5% agarose gel and purified using a QIAquick gel extraction kit (QIAGEN Inc.) according to the manufacturer’s instructions. The gel-purified PCR products were purified again using 1 volume of SPRI beads and eluted in 20 μl water. The SPRI-purified sequencing libraries were quantified with a 4200 TapeStation System (Agilent Technologies) using a D1000 ScreenTape assay and subjected to next-generation sequencing on the Illumina NovaSeq platform.

### NGS data processing

Raw sequencing reads obtained from sequencing the V_H_-CH1 region of B cells in peripheral blood were processed using a custom pipeline. The pipeline included adapter trimming and quality filtering; unique molecular identifier (UMI) processing; V(D)J gene annotation; clustering; quality control; and diversity analysis.

The forward reads (R1) and reverse reads (R2) of the raw NGS data were merged using paired-end read merger (PEAR) v0.9.10 with the default settings^32^. The merged reads were q-filtered under the q20p95 condition, resulting in 95% of base pairs in the reads having a Phred score greater than 20. Primer positions were identified in the quality-filtered reads, and primer regions were trimmed to remove the effects of primer synthesis errors while allowing one substitution or deletion.

Based on the primer recognition results, UMI sequences were extracted, and reads were clustered according to the UMI sequences. To eliminate index misassignment, we subclustered the clustered reads based on the similarity of the reads (allowing 5 mismatches in each subcluster) and matched the majority subcluster to the UMI. The subclustered reads were aligned using the multiple sequence alignment tool Clustal Omega v1.2.4 with the default settings^33, 34^. Consensus calling was performed by selecting major frequency bases at every position of the aligned sequences. The number of reads in the consensus sequence was redefined as the number of UMI subclusters belonging to the consensus sequence.

Sequence annotation consisted of isotype annotation and V(D)J annotation. The consensus sequence was divided into a V(D)J region and a constant region. The isotype of the consensus sequence was annotated by aligning the extracted constant region with the constant gene of the International Immunogenetics Information System (IMGT) ^35^. Then, the V(D)J region of the consensus sequence was annotated using an updated version of IgBLAST (v1.17.1) ^36^. Among the annotation results, the IGHV genes, IGHJ genes, HCDR3 sequences, and number of SHMs were extracted for further analysis. The number of SHMs was counted only in the aligned IGHV genes. The nonfunctional consensus reads were defined and filtered using the following criteria: (i) sequence length shorter than 250 base pairs, (ii) presence of a stop codon or frameshift in the entire amino acid sequence, (iii) failure to annotate one or more HCDR1, HCDR2 and HCDR3 regions, and (iv) failure to annotate the isotype.

### Construction of scFv phage display library

Six human scFv phage display libraries were constructed using the total RNA prepared from the sixth blood sample for vaccine Nos. 22, 27, 32, 35, 39 and 43 as described previously^23^. Briefly, for the V_H_ and V_κ_/V_λ_ genes, total RNA was employed to synthesize cDNA using the SuperScript IV First-Strand Synthesis System (Invitrogen) with gene-specific primers targeting the J_H_ and C_κ/λ_ genes (primer No.17–22), respectively. After cDNA synthesis, 1.8 volumes of SPRI beads (Beckman Coulter) were used to purify cDNA, which was eluted in 20 μl of water. The purified cDNA (11.25 μl) of the V_H_ gene was subjected to second-strand synthesis in a 25 μl reaction volume using IGHV gene-specific primers (primer No. 23–32, 7.5 μl) and a KAPA Biosystems kit (Roche). The reaction conditions were as follows: 95°C for 3 min, 98°C for 1 min, 60°C for 1 min, and 72°C for 5 min. In the case of the V_κ_/V_λ_ gene, the eluted cDNA (17.25 μl) was used for the first round of PCR synthesis in a 25 μl reaction volume using V_κ_/V_λ_ and J _κ_/J _λ_ gene-specific primers (primer No. 33–69, 0.75 μl). The PCR conditions were as follows: 95°C for 3 min; 4 cycles of 98°C for 1 min, 60°C for 1 min, and 72°C for 1 min; and 72°C for 10 min. After second-strand synthesis for the V_H_ gene or PCR amplification of the V_κ_/V_λ_ gene, double-stranded DNA (dsDNA) was purified with 1 volume of SPRI beads and eluted in 40 μl of water. V_H_ and V_κ_/V_λ_ genes were amplified using 10 μl of purified dsDNA, 2.5 pmol of the primers (primer No. 70–73), and KAPA Biosystems kit components in a 50 μl total reaction volume with the following thermal cycling program: 95°C for 3 min; 30 cycles of 98°C for 30 s, 60°C for 30 s, and 72°C for 1 min; and 72°C for 10 min. Then, the amplified V_H_ and V_κ_/V_λ_ genes were subjected to electrophoresis on a 1.5% agarose gel and purified using a QIAquick gel extraction kit (QIAGEN Inc.) according to the manufacturer’s instructions. The purified V_H_ and V_κ_/V_λ_ gene fragments (100 ng) were mixed and subjected to overlap extension PCR to generate scFv genes using 2.5 pmol of the overlap extension primers (primer No. 74 and 75) using the KAPA Biosystems kit. The PCR conditions were as follows: 95°C for 3 min; 25 cycles of 98°C for 20 s, 65°C for 15 s, and 72°C for 1 min; and 72°C for 10 min. The amplified scFv gene was purified and cloned into a phagemid vector as described previously^37^.

### Selection of BA.1RBD-binding clones

Human scFv phage display libraries with 7.1 × 10^8^, 6.1 × 10^8^, 8.1 × 10^8^, 8.3 × 10^8^, 7.9 × 10^8^, and 7.4 × 10^8^ colony-forming units were generated using cDNA prepared from vaccinee Nos. 22, 27, 32, 35, 39 and 43, respectively. The libraries were subjected to five rounds of biopanning against the recombinant SARS-CoV-2 BA.1 RBD protein (Sino Biological Inc.) as described previously^38^. Immune tubes (SPL, 43015) coated with 17 μg of the BA.1 RBD were used for the first round, and magnetic beads (Invitrogen, Dynabeads M-270 epoxy) conjugated with 1.4 μg of the BA.1 RBD protein were used for the other rounds. After each round of biopanning, the bound phages were eluted and amplified for the next round of biopanning. For the selection of BA.1 RBD-reactive scFv phage clones, individual phage clones were amplified from the titration plate of the last round and subjected to phage ELISA as described previously^39^. The genes encoding BA.1 RBD-reactive scFv clones were identified using phagemid DNA prepared from phage clones and Sanger nucleotide sequencing as described previously^38^. A recombinant scFv protein fused with human IgG1 FC and the HA peptide (scFv-hFc-HA) was expressed using a mammalian expression system and purified as described previously^39^.

### HCDR3-randomized scFv phage display library

The gene fragment encoding the V_H_ region with a randomized HCDR3 sequence and another gene fragment encoding the rest of the scFv were amplified using phagemid DNA of clone 27-60, primers for randomization (primer No.76-79) and the KAPA Biosystems kit. The PCR conditions were as follows: 95°C for 3 min; 25 cycles of 98°C for 20 s, 65°C for 15 s, and 72°C for 1 min; and 72°C for 10 min. Then, the amplified genes were subjected to electrophoresis on a 1.5% agarose gel and purified using a QIAquick gel extraction kit (QIAGEN Inc.) according to the manufacturer’s instructions. The purified gene fragments (200 ng) were mixed and subjected to overlap extension PCR to generate scFv genes using 2.5 pmol of the overlap extension primers (primer No.76 and 79) and the KAPA Biosystems kit. The PCR conditions were as follows: 95°C for 3 min; 25 cycles of 98°C for 20 s, 65°C for 15 s, and 72°C for 1 min; and 72°C for 5 min. The amplified scFv genes were purified and cloned into a phagemid vector as described previously^37^. For phage ELISA, individual phage clones were amplified from the titration plate and subjected to phage ELISA as described previously^40^.

### Microneutralization assay

The ancestral SARS-CoV-2 (βCoV/Korea/KCDC03/2020 NCCP43326), Alpha B.1.1.7 (hCoV-19/Korea/KDCA51463/2021 (NCCP 43381), Beta B.1.351 (hCoV-19/Korea/KDCA55905/2021 (NCCP 43382), Gamma P.1 (hCoV-19/Korea/KDCA95637/2021 (NCCP 43388), Delta B.1.617.2 (hCoV-19/Korea/KDCA119861/2021 (NCCP 43390), and Omicron B.1.1.529 (hCoV-19/Korea/KDCA447321/2021 NCCP43408) viruses were obtained from the Korea Disease Control and Prevention Agency. The viruses were propagated in Vero cells (ATCC, CCL-81) in Dulbecco’s Modified Eagle’s Medium (DMEM, Welgene) in the presence of 2% fetal bovine serum (Gibco, Thermo Fisher Scientific Inc.), as described previously^23, 41^.

Neutralization assays were performed as described previously^23, 42^. Briefly, Vero cells were seeded in 96-well plates (1.5×10^4^ or 0.5×10^4^ cells/well) in Opti-PRO SFM (Thermo Fisher Scientific Inc.) supplemented with 4 mM L-glutamine and 1× Antibiotics-Antimycotic (Thermo Fisher Scientific Inc/) and grown for 24 h at 37°C in a 5% CO_2_ environment. Recombinant scFv-hFc proteins were diluted from 100 to 0.1953 µg/ml (twofold) in phosphate-buffered saline (PBS, Welgene) and mixed with 100 or 500 TCID_50_ of SARS-CoV-2. Then, the mixture was incubated for 30 min at 37°C and added to the cells in tetrads, followed by incubation for 4 or 6 days at 37°C in a 5% CO_2_ environment. The cytopathic effect (CPE) in each well was visualized following crystal violet staining 4 or 6 days post infection. The IC_50_ values were calculated using the dose-response inhibition equation of GraphPad Prism 6 (GraphPad Software).

### Construction of BCR HC clonotype-3^rd^

The most frequent BCR HC sequences of the 27-60, 35-15, 35-46, 43-09, and 43-34 clonotypes in the third BCR repertoires were mapped to the chronological BCR repertoires following the same definition of BCR HC clonotypes. All the mapped nucleotide sequences were aligned using the multiple sequence alignment tool Clustal Omega v1.2.4 with the default settings^33, 34^. The aligned mapped nucleotide sequences were interpreted using a phylogenetic tree generated by IgPhyML v1.1.3 052020 using the HLP model option^43^. The phylogenetic trees were plotted using Interactive Tree of Life (iTOL) ^44^.

## Data availability

All sequencing data are available from the National Center for Biotechnology Information (www.ncbi.nlm.nih.gov/) under accession number PRJNA945512 (SRA).

## Code availability

NGS data preprocessing was performed as described in a previous study (Kim, S. I. *et al*. Sci Transl Med 13 (2021)). The code use in this study is available upon request for academic research purposes.

## Acknowledgements

This research was supported by the Korea Dementia Research Center (KDRC, HU20C0339), Korea Health Technology R&D Project through the Korea Health Industry Development Institute (KHIDI, HI23C0521), the BK21 FOUR program, the National Research Foundation of Korea (NRF-2020R1A3B3079653 NRF-2020M3H1A1073304, NRF-2017M3A9G6068245, NRF-2022M3A9J1081343 and NRF-2023M3A9G6057281), the SNUH Research Fund (03-2021-0080), and the Inter-university Semiconductor Research Center. The pathogen resources (NCCP43326, NCCP43381, NCCP43382, NCCP43388, NCCP43390, NCCP43408) for this study were provided by the National Culture Collection for Pathogens. J. Choi is grateful for financial support from the Hyundai Motor Chung Mong-Koo Foundation.

## Author contributions

S.P. designed and conducted all experiments, performed analyses, interpreted experimental results, and wrote and revised the paper. J. Choi performed the bioinformatic analyses, visualized and interpreted the results, and wrote and revised the paper. Y.L., J.N. and N.P.K. performed the bioinformatic analysis. J.L. conducted experiments related to the microneutralization assay. D.Y. conducted experiments related to the construction of repertoire libraries for NGS. G.C. and S.J.K. contributed to experiments related to the expression of recombinant proteins. K.K., P.C., N.J.K., and W.P. enrolled the study participants and collected the study samples. M.O. conceived and designed the study, and interpreted all results. S.T.K. supervised all the experiments with infectious viruses and microneutralization assay. S. Kwon conceived the study and designed and supervised the bioinformatics analysis. J. Chung conceived the study, designed and supervised all experiments, interpreted all results, and wrote the paper.

## Additional information

Supplementary Information is available for this paper

Correspondence and requests for materials should be addressed to J. Chung.

Reprints and permissions information is available at www.nature.com/reprints.

## Extended data

**Extended Data Fig. 1.**
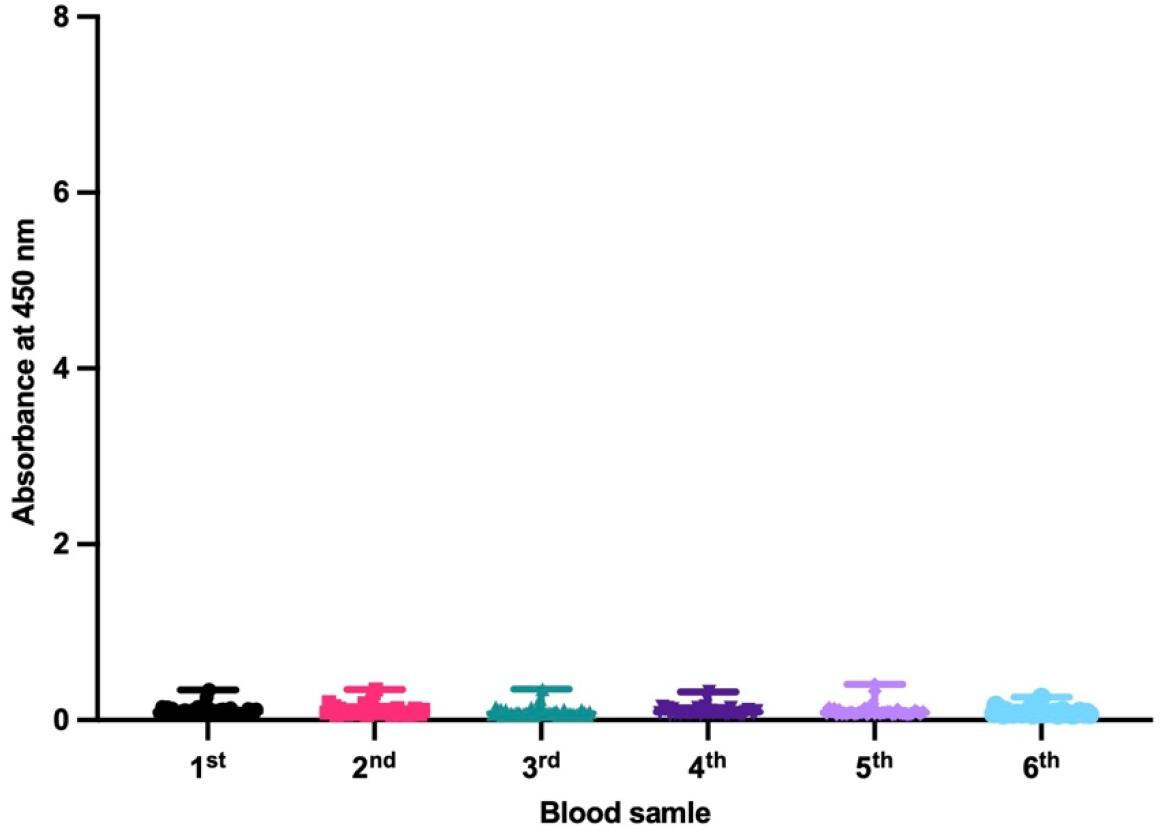
Plasma levels of antibodies against the SARS-CoV-2 N protein. Reactivity of plasma from 41 vaccinees to the SARS-CoV-2 N protein (Sino Biological) was tested by ELISA. The wells of microtiter plates were coated with the SARS-CoV-2 N protein and blocked. After washing, plasma samples diluted in blocking buffer (1:2,500-fold) were added to wells. The amount of bound antibody was determined using a horseradish peroxidase (HRP)-conjugated rabbit anti-human IgG antibody (Invitrogen).

**Extended Data Fig. 2.**
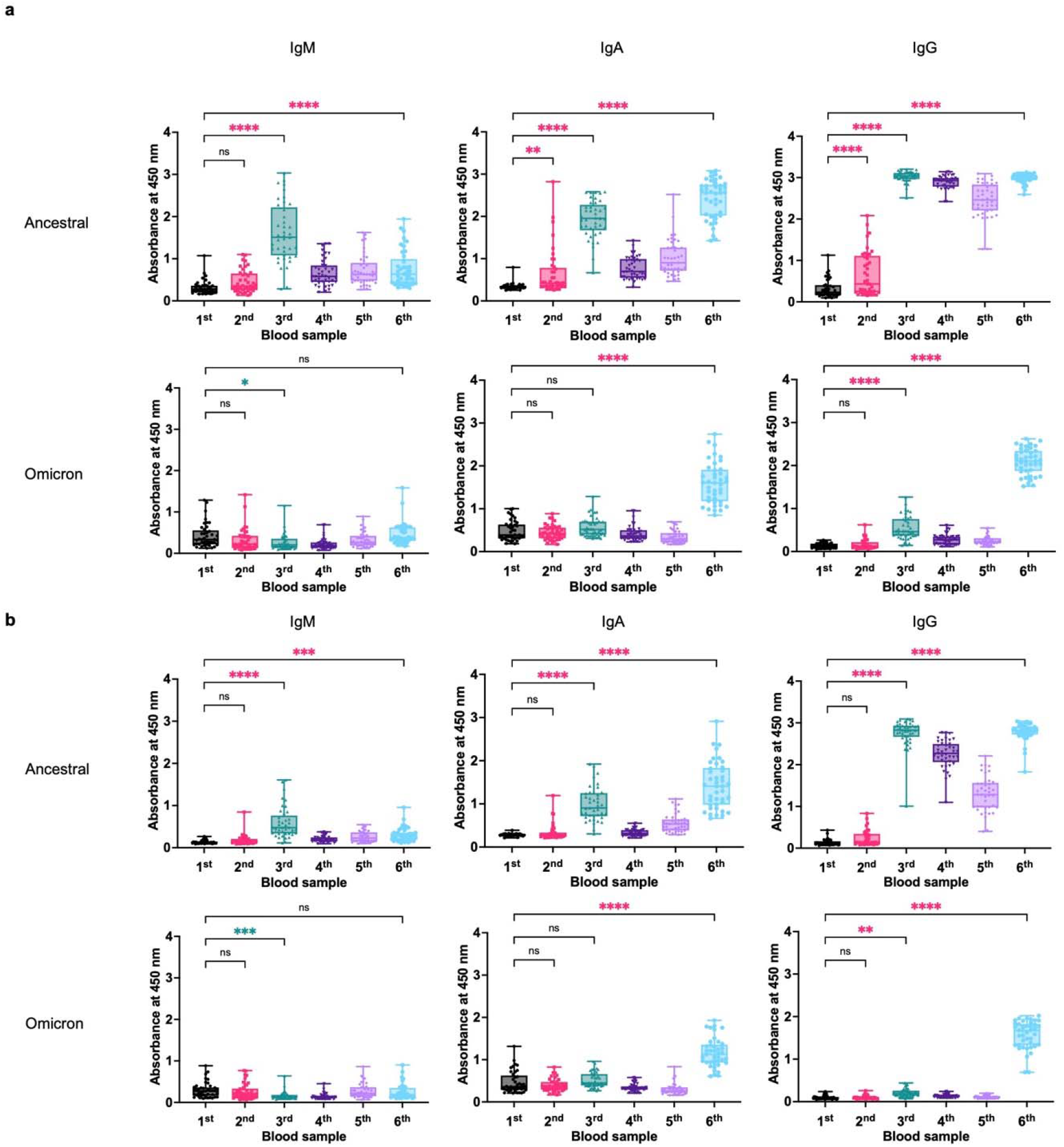
Plasma levels of antibodies against ancestral and BA.1 RBD. Plasma levels of antibodies to SARS-CoV-2 ancestral and BA.1 RBDs were measured in 41 vaccinees after (**a**) 100-fold or (**b**) 500-fold dilution. All experiments were performed in duplicate, and the average value for each vaccinee was plotted. The P value was calculated using one-way ANOVA. ns, not significant, p > 0.05; **, p < 0.01; ***, p < 0.001; ****, p < 0.0001. The increases and decreases in antibody levels are marked with red and green asterisks, respectively

**Extended Data Fig. 3.**
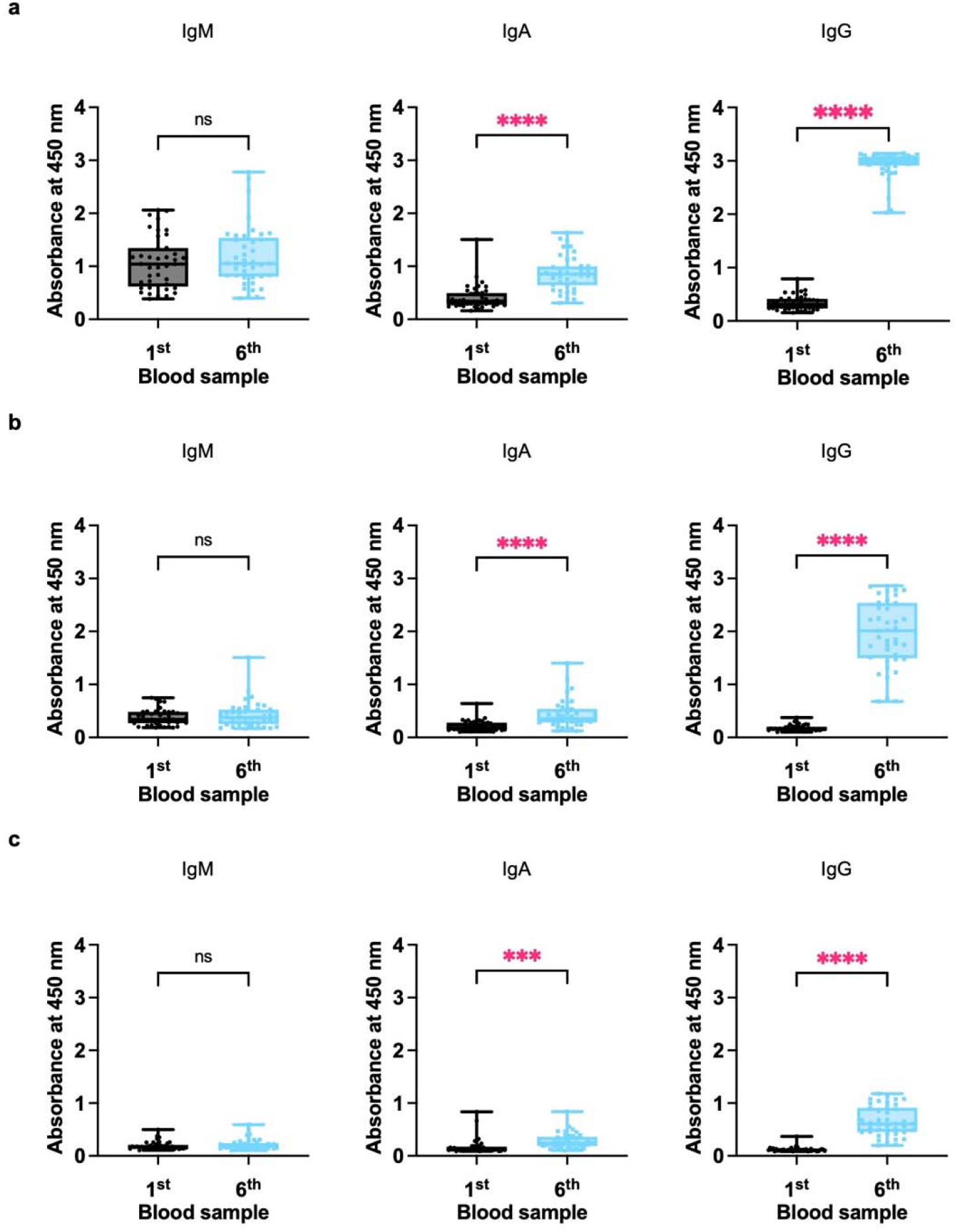
Plasma levels of antibodies against the RBD of Omicron subvariant BQ.1.1. Plasma levels of antibodies to the Omicron subvariant BQ.1.1. RBD (Sino Biological) were determined after (**a**) 100-, (**b**) 500- or (**c**) 2,500-fold dilution. All experiments were performed in duplicate, and the average value for each vaccinee was plotted. The P value was calculated using one-way ANOVA. ns, not significant, p > 0.05; ***, p < 0.001; ****, p < 0.0001.

**Extended Data Fig. 4.**
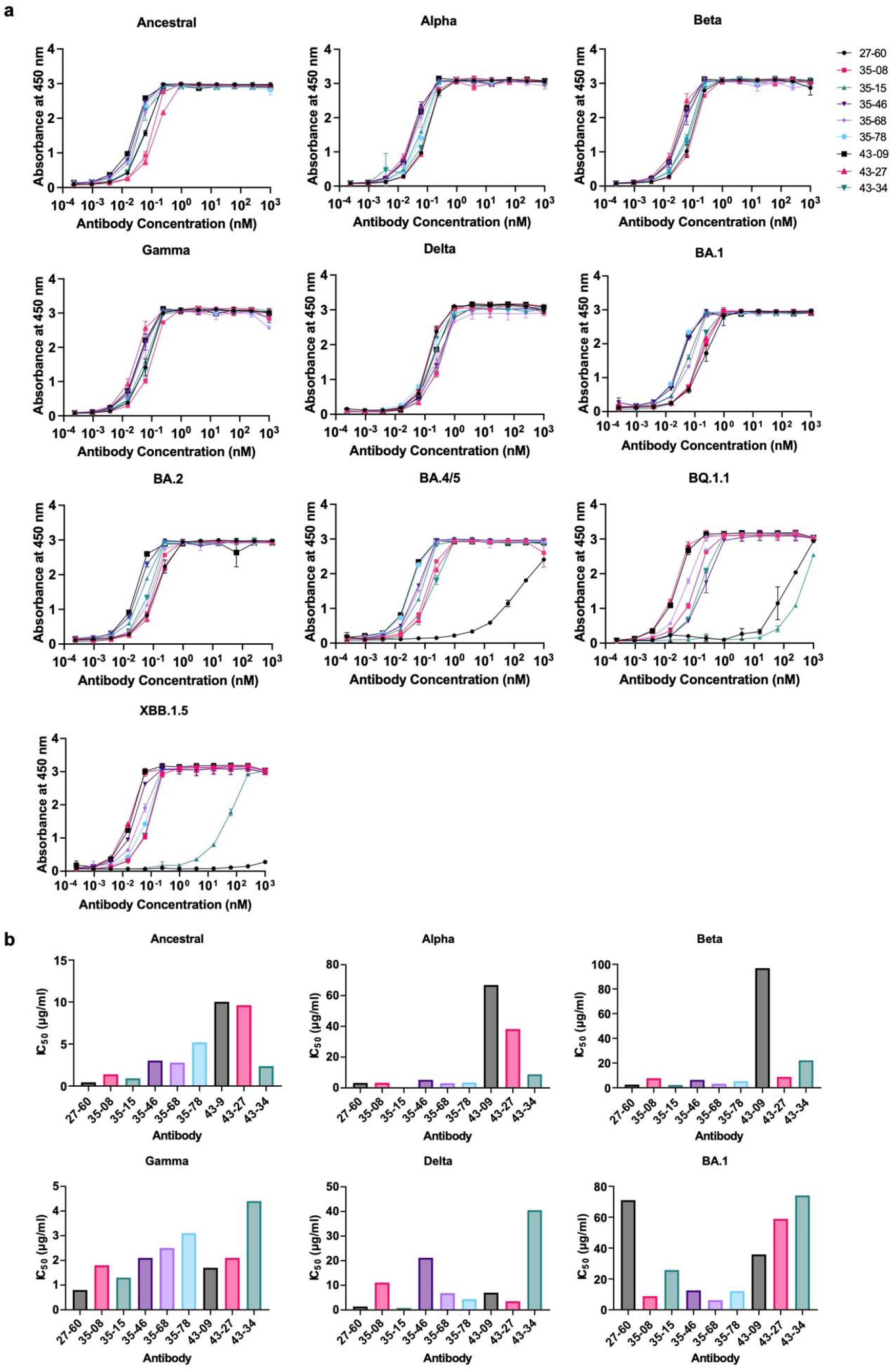
Characteristics of BA.1 RBD-binding scFv clones. **a,** Reactivity of BA.1 RBD-binding scFv clones from the phage display library was tested against SARS-CoV-2 ancestral, Alpha, Beta, Gamma, Delta, and Omicron-sublineage (BA.1, BA.2, BA.4/5, BQ.1.1 and XBB.1.5) RBD proteins in the form of a recombinant scFv-hFc-HA protein. **b,** The neutralization efficacy of BA.1 RBD-reactive scFv clones was tested in a microneutralization assay using ancestral SARS-CoV-2 (βCoV/Korea/KCDC03/2020 NCCP43326), Alpha B.1.1.7 (hCoV-19/Korea/KDCA51463/2021 (NCCP 43381), Beta B.1.351 (hCoV-19/Korea/KDCA55905/2021 (NCCP 43382), Gamma P.1 (hCoV-19/Korea/KDCA95637/2021 (NCCP 43388), Delta B.1.617.2 (hCoV-19/Korea/KDCA119861/2021 (NCCP 43390), and Omicron B.1.1.529 (hCoV-19/Korea/KDCA447321/2021 NCCP43408) strains.

**Extended Data Fig. 5.**
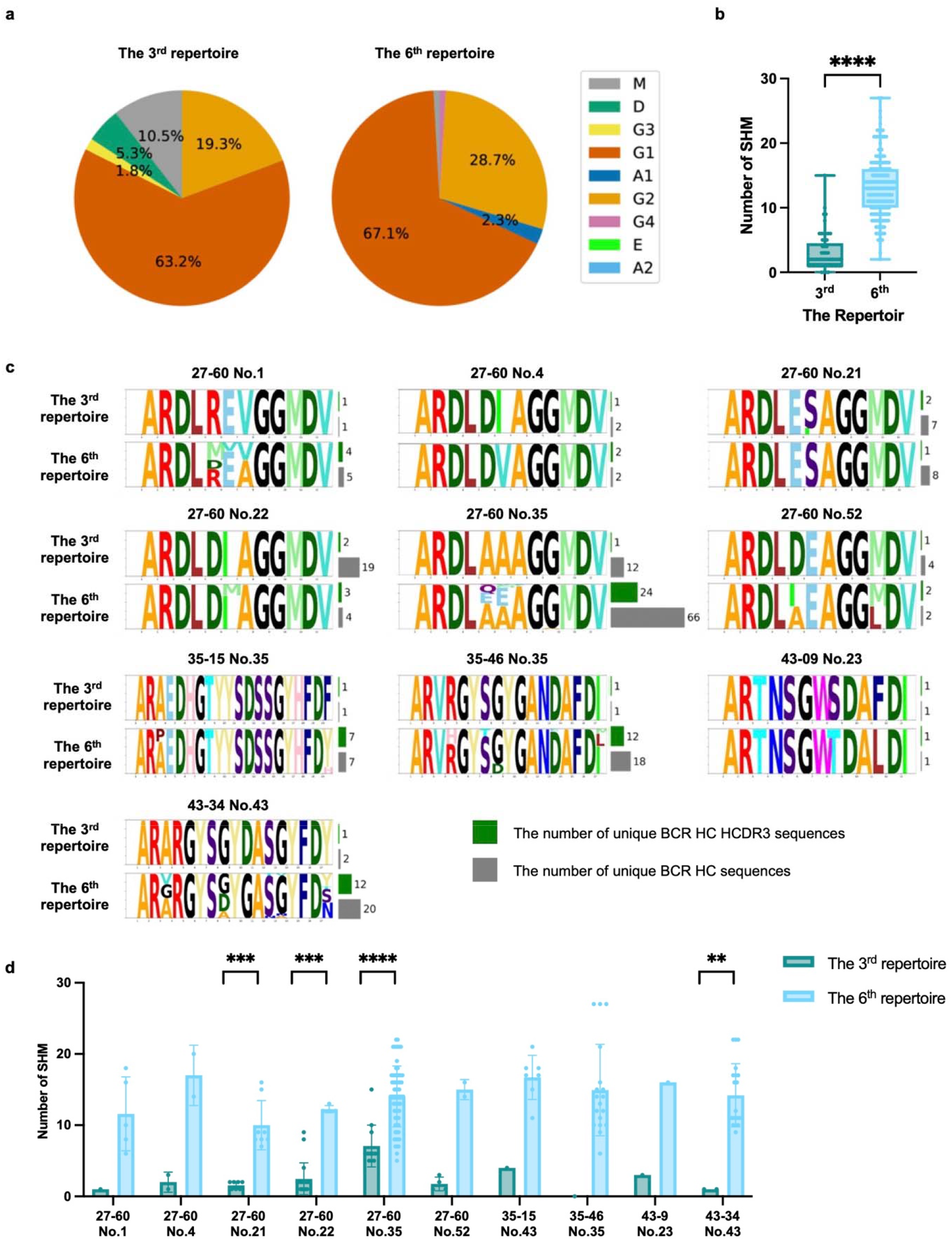
The characteristics of the BCR HC sequences of nine Omicron RBD-reactive clonotypes mapped to the third and sixth repertoires. **a,** The isotype compositions of the BCR HC sequences in the third and sixth repertoires (n = 57 and 216, respectively). **b,** Numbers of SHMs in BCR HC sequences at the third and sixth repertoires (n = 57 and 216, respectively). The P value was calculated using a two-tailed independent t test. ****, p < 0.0001. **c,** Diversification of BCR HC sequences occurred from the third to the sixth repertoire through SHM and HCDR3 sequence variation. The numbers of unique BCR HC sequences and HCDR3 sequences are counted at the nucleotide level. **d,** Numbers of SHMs in the BCR HC sequences of clonotypes found in the third and sixth repertoires of the same vaccinee. The P value was calculated using the Mann-Whitney U test. In the case of repertoires for which statistical significance is not indicated, statistical significance was not calculated, as the number of clones included in the third or sixth repertoire was only one or two. ns, not significant, p > 0.05; *, p < 0.05; **, p < 0.01; ***, p < 0.001; ****, p < 0.0001.

**Extended Data Fig. 6.**
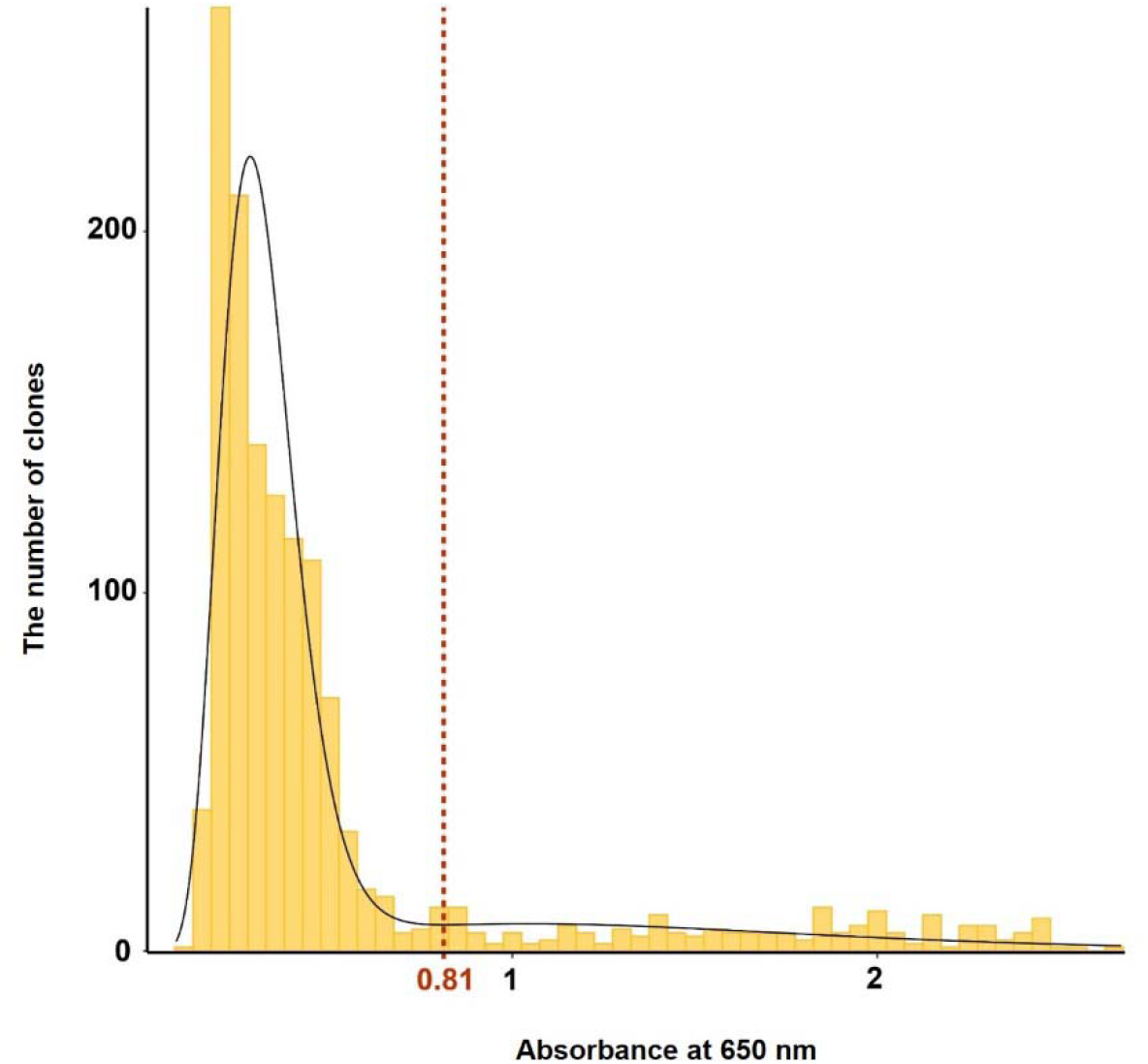
Determination of the threshold for RBD-reactive clones in the HCDR3-randomized scFv phage display library. After phage ELISA using both ancestral and BA.1 RBDs, the absorbance of each well was determined. The absorbance value was approximated with a Gaussian distribution by the binned kernel estimation method under the Gaussian kernel setting (black line). The chosen threshold absorbance value for a positive clone was 0.81 (red vertical line), which corresponded to the valley point in the distribution plot.

**Extended Data Fig. 7.**
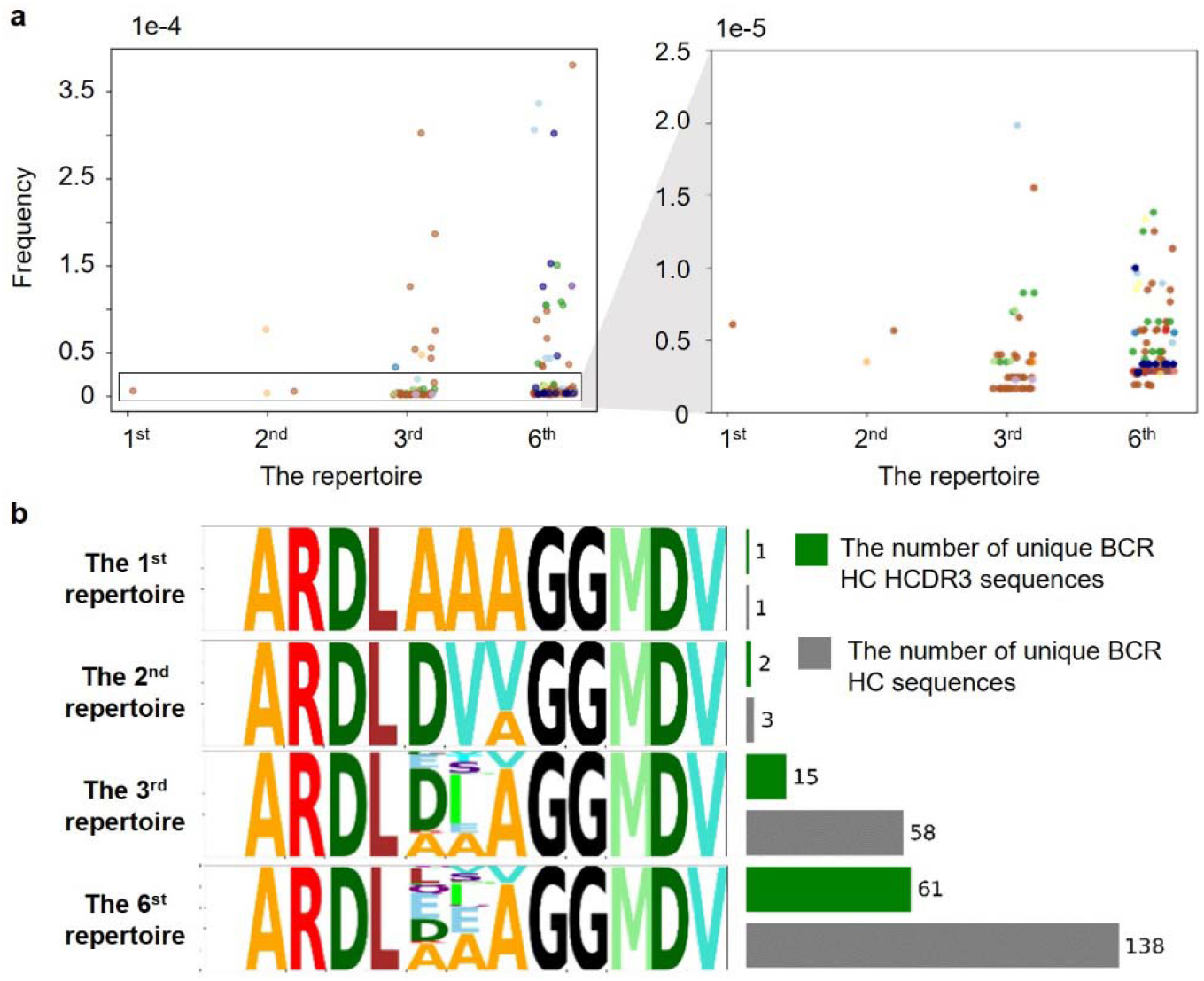
The frequency distribution and sequence diversity of BCR-HC XXA/V clonotypes after the first, second, and third injections. **a,** The frequency distribution of BCR HC XXA/V clonotypes. The BCR HC XXA/V clonotypes with the same HCDR3 amino acid sequence are plotted in the same color. **b,** Diversity of BCR HC XXA/V sequences in individual repertoires. The numbers of unique BCR HC sequences and HCDR3 sequences are counted at the nucleotide level.

**Extended Data Fig. 8.**
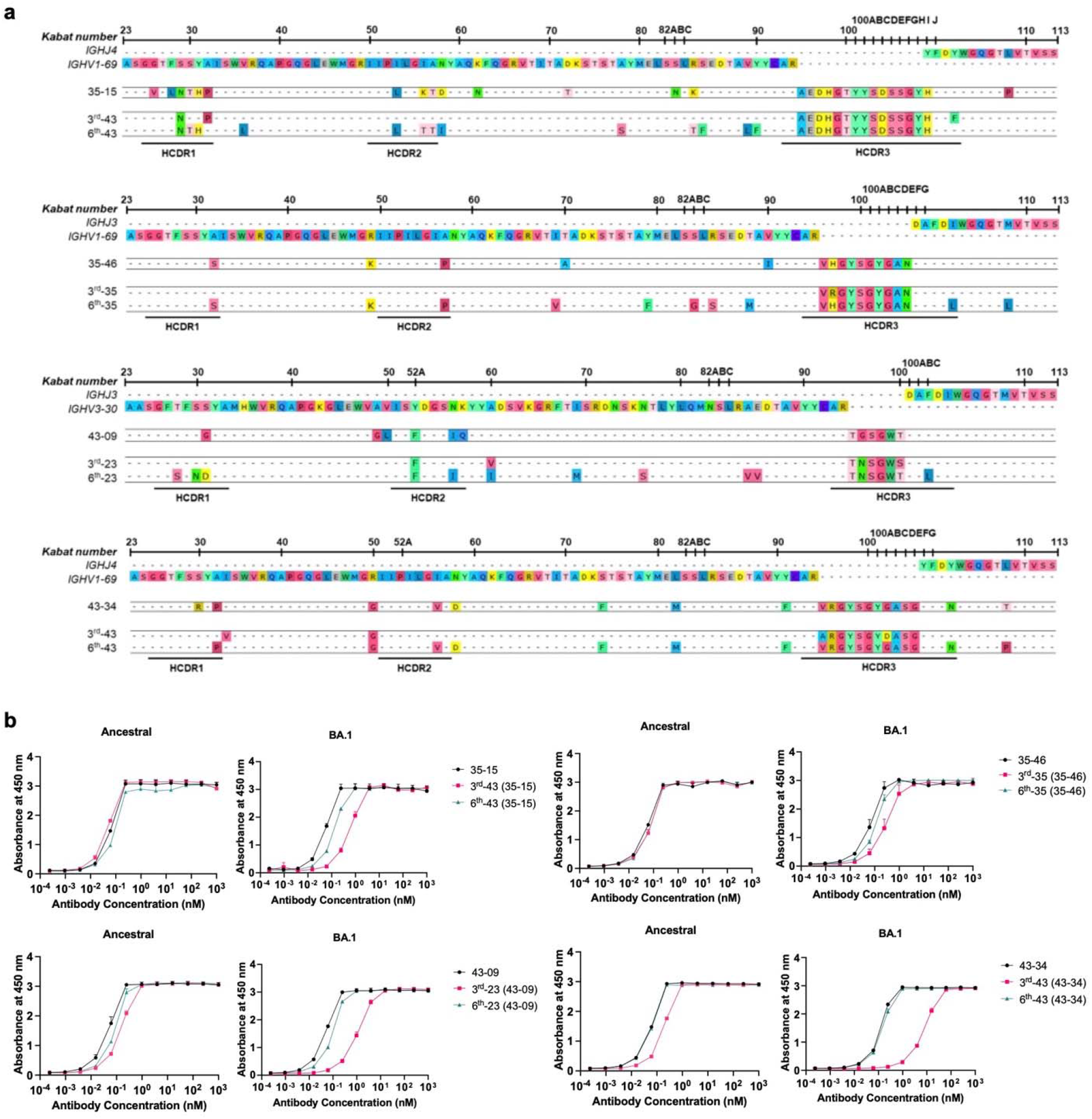
**a,** BCR HC sequences of four clonotypes found in vaccinees in their third and sixth BCR repertoires with the highest frequency. **b,** Reactivity of recombinant scFv-hFc-HA proteins encoded by individual BCR HC sequences and the light chain gene of each scFv clone to the ancestral and BA.1 RBDs.

**Extended Data Fig. 9.**
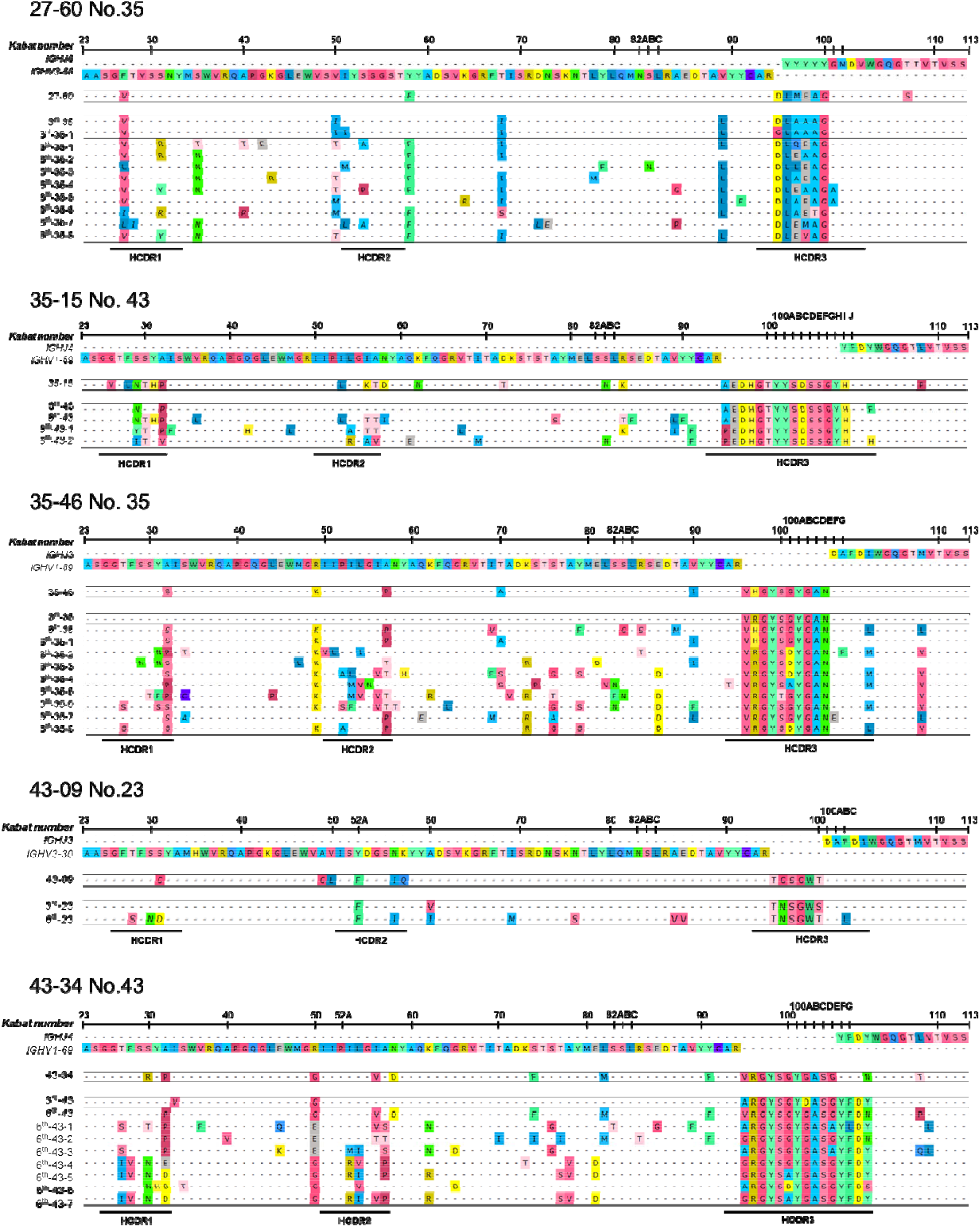
The BCR HC sequences of the 27-60, 35-15, 35-46, 43-09, and 43-34 clonotypes-3^rd^ found in vaccinees 35, 43, 35, 23, and 43.

**Extended Data Fig. 10.**
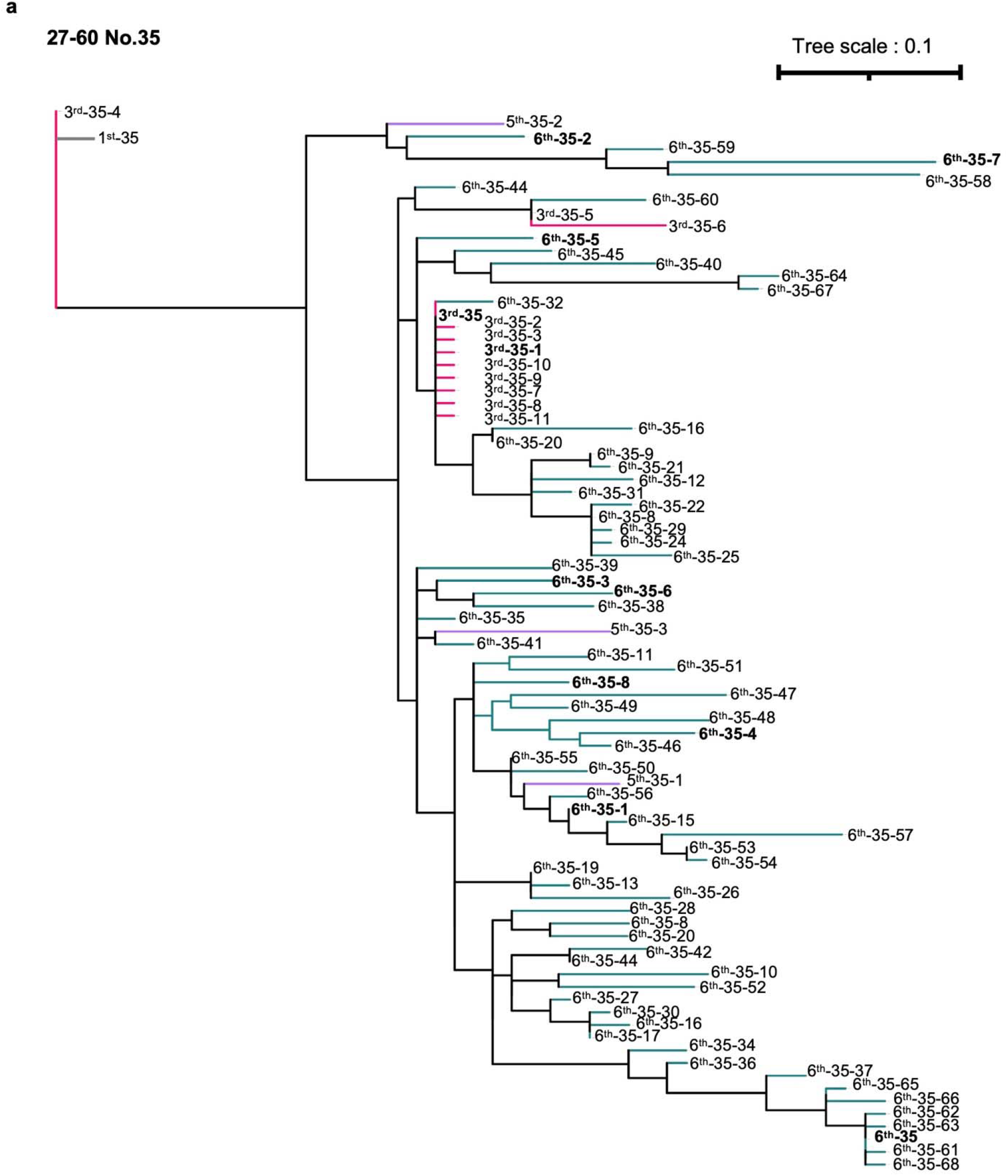
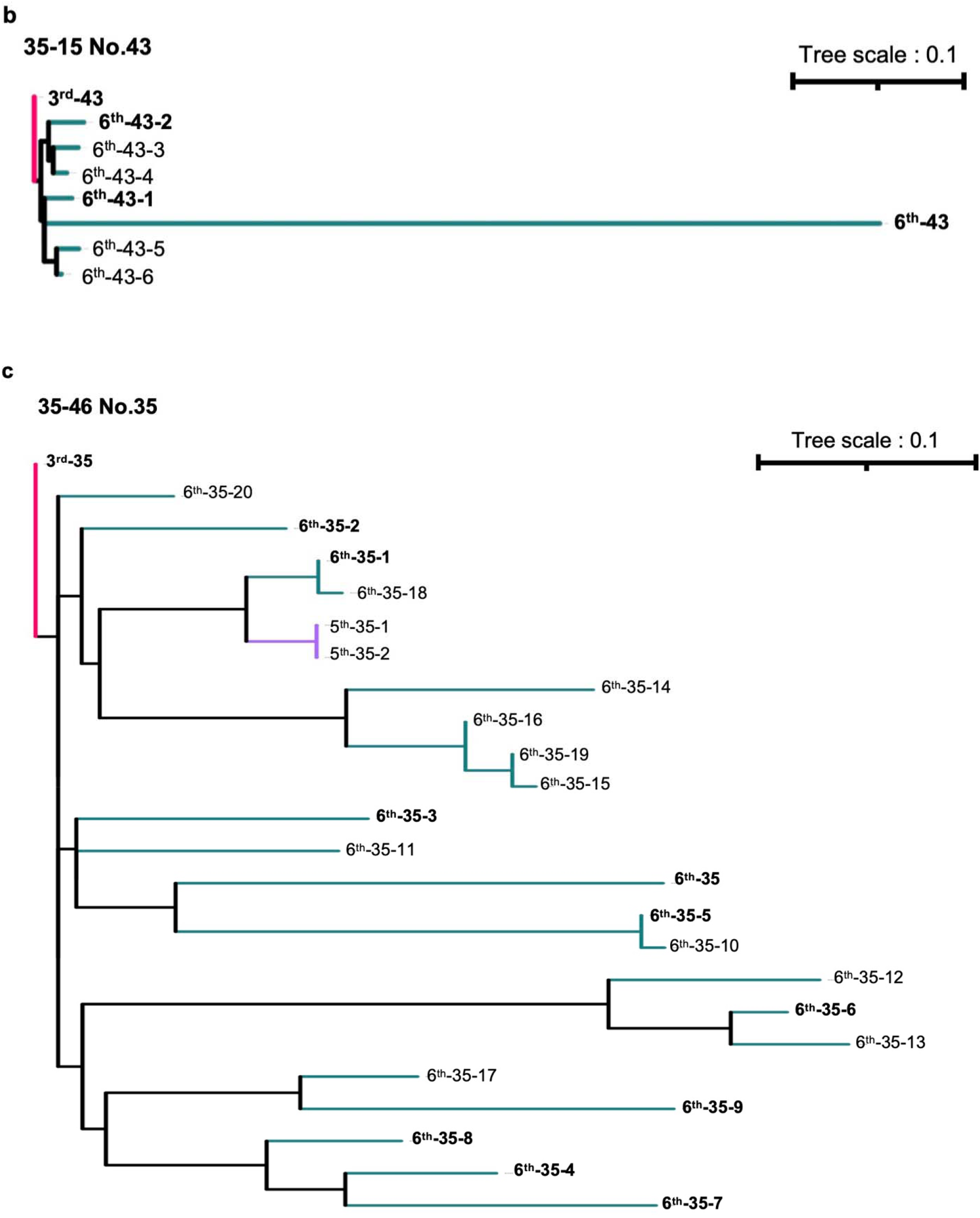
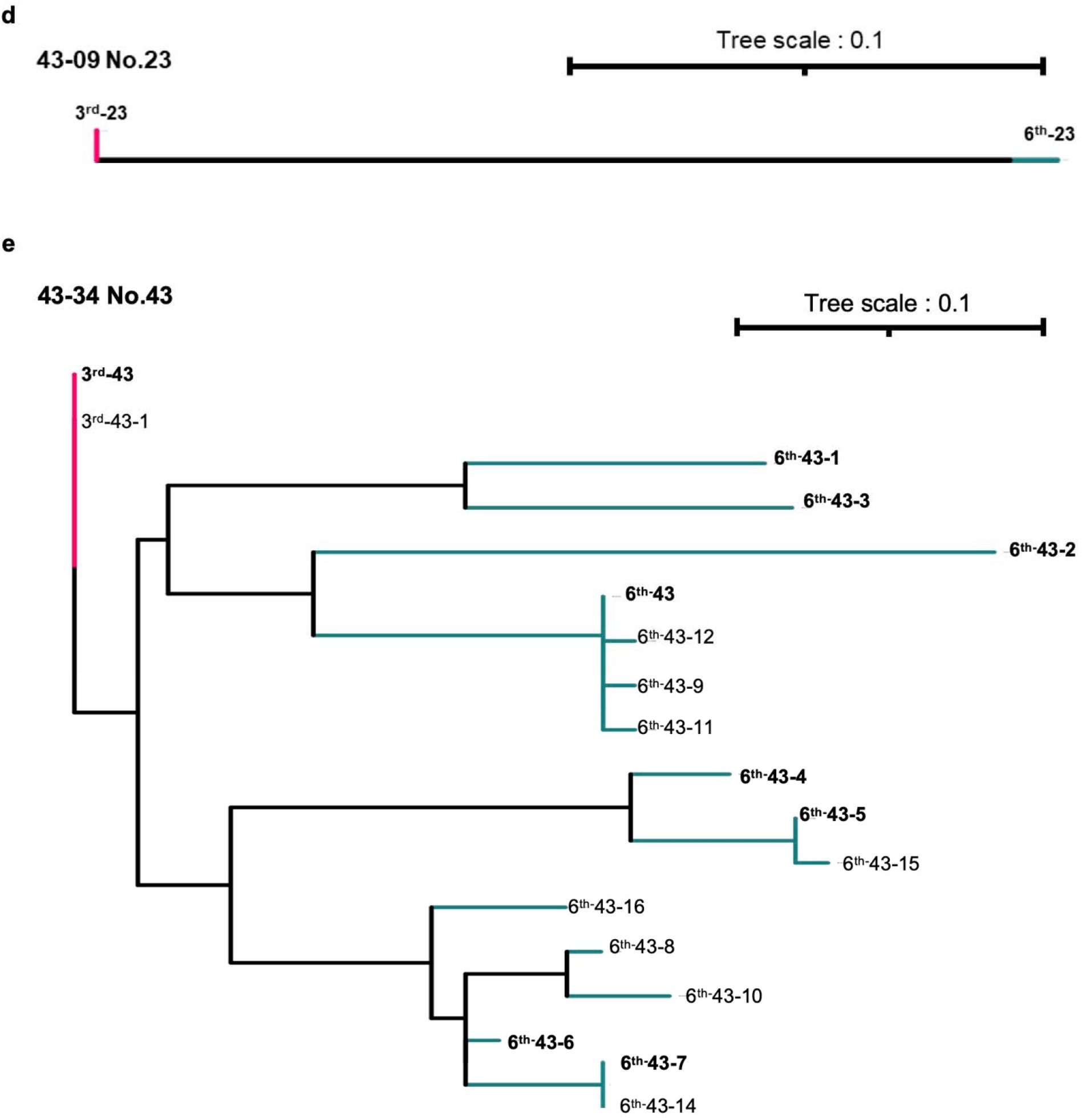
Phylogenetic trees of the 27-60, 35-15, 35-46, 43-09, and 43-34 clonotypes-3^rd^ found in vaccinees 35, 43, 35, 23, and 43. The BCR sequences that were synthesized to confirm the reactivity to ancestral RBD and those of Omicron subvariants are shown in bold.

